# Deep learning-based detector of invasive alien frogs, *Polypedates leucomystax and Rhinella marina*, on an island at invasion front

**DOI:** 10.1101/2024.07.27.605431

**Authors:** Kaede Kimura, Ibuki Fukuyama, Kinji Fukuyama

## Abstract

This study aimed to develop and evaluate a deep learning-based detector for Southeast Asian treefrog (*Polypedates leucomystax*) and cane toad (*Rhinella marina*) on Iriomote Island, a Natural World Heritage Site located 30 km from established populations of these alien species on the nearby Ishigaki Island. Although a deep learning model typically requires local training data to be most accurate, the alien frogs have been eradicated from Iriomote Island, making such data unavailable. To address the data gap, we first trained the BirdNET model with acoustic data collected on Ishigaki Island, where these species were common, as well as native frog calls from Iriomote Island. Next, we evaluated model performance using (1) wild sounds on Ishigaki Island, and (2) sounds obtained by playing back the calls of the alien species on Iriomote Island. Model precision and recall for wild sounds were both 0.987 for *P. leucomystax* and 1 for *R. marina*. For playback sounds, model recall values decreased (0.629 for *P. leucomystax* and 0.906 for *R. marina*), while precisions remained nearly identical (1 for both *P. leucomystax* and *R. marina*). Despite the lower recall particularly for *P. leucomystax*, playback survey dates were mostly identifiable from the high number of detections. These results suggest that data from Ishigaki Island enabled training a model with adequate, though not complete, generalization across invasion front.

## Introduction

Biological invasion is a major threat to biodiversity worldwide, and eradication of introduced species is most feasible during the early phase of invasion (IPBES 2023). Early detection of invasive alien species has been facilitated by novel monitoring technologies like remote sensing, environmental DNA (eDNA), and machine learning (Fricke and Olden 2023). Deep learning, a form of machine learning, is gaining attention for automated biological monitoring because of its high accuracy and scalability (Norouzzadeh et al. 2018; Kahl et al. 2021). Recently, several studies attempted to detect invasive alien species in the wild with deep learning. Examples include detecting *Anolis* lizards (Aota et al. 2021), lionfish (Martínez-González et al. 2021), and Asian black hornets (O’Shea-Wheller et al. 2024) from images, and barred owls (Kelly et al. 2023) and bullfrogs (Bota et al. 2024) from sounds. These studies suggest that a deep learning model can detect invasive alien species when local training samples (i.e., samples collected at sites of actual application) are available.

However, it is impossible to acquire local training data when the purpose of a project is preparedness at an invasion front, and the model trained on *ex situ* data can have lower performance due to domain shift. Domain shift refers to the situation where training and target data have different characteristics, often reducing model accuracy. For instance, models trained with camera traps at specific locations perform poorly at new locations because of differences in background and lighting conditions (Beery et al. 2018; Schneider et al. 2020; Norman et al. 2023). At the front of biological invasion, early detection systems are deployed in locations where training data of the target alien species are unavailable (e.g., Taylor et al. 2017). The expected accuracy reduction of the model trained on *ex situ* data, as well as whether the magnitude of accuracy reduction matters in a real-world setting, have rarely been examined, though this situation would be common.

Iriomote Island, a tropical island designated as a Natural World Heritage Site in Japan, has been threatened by the invasion of two alien frog species, cane toad (*Rhinella marina*) and Southeast Asian treefrog (*Polypedates leucomystax*) (National Institute for Environmental Studies 2024). *Rhinella marina* is not only a voracious predator but also a toxic prey, and especially the latter makes this species highly invasive to the local biodiversity (Shine 2010). The ecological impacts of *P. leucomystax* have not been studied well, but possible negative effects on native anurans through resource competition, reproductive interference, and parasite transfer are concerning (Ota et al. 2004; National Institute for Environmental Studies 2024). These two species have established dense populations at the nearby Ishigaki Island (about 30 km apart) and been unintentionally transported to Iriomote Island for dozens of times. Continuous detection and eradication efforts have prevented these species from establishing and spreading on Iriomote Island to date (Nakajima et al. 2005), but constraints on resources and manpower limit surveillance frequency and coverage. Therefore, developing a system that can automatically detect the invasive alien frogs on Iriomote Island are needed.

Our objectives in this study were to develop and test an early detection system for *R. marina* and *P. leucomystax* on Iriomote Island, where local training data are unavailable. First, we trained a deep learning model to identify the two alien and six native frog species from their calls, as frog populations can be effectively monitored through sounds (Dorcas et al. 2009). During this process, the training data of the alien species was obtained from the nearby Ishigaki Island. We expected minimal domain shift in acoustic data between Iriomote Island and Ishigaki Island due to their faunal similarity (e.g., all amphibian species are shared; Matsui and Maeda 2018). Next, we evaluated the model performance in detecting alien frogs on both Ishigaki and Iriomote Islands. On Iriomote Island, where these alien species had no established population, we tested the model using field playback of their calls. Third, we assessed the similarity between playback sounds and wild calls by mapping acoustic features (i.e., model embeddings). Finally, to assess the seasons for effective acoustic monitoring, we investigated breeding phenology of *P. leucomystax* and *R. marina* by applying the trained model to the one-year acoustic data from Ishigaki Island. In doing so, we aimed to obtain a deep learning model that generalizes well across the invasion front for real-world deployment.

## Materials and methods

### Acoustic data collection

Acoustic data were collected from three sites on Ishigaki Island and three sites on Iriomote Island (Fig. 1). We selected habitats including roadside ponds (sites 1, 3, 4), mixture of sugar cane field and rice paddy (site 2), artificial concrete pond (site 3), rice paddy (site 5), and roadside ditch (site 6), since *P. leucomystax* and *R. marina* commonly found in disturbed environments on these islands. We placed acoustic recorders (RR-XS455, Panasonic) at sites 1–4 from March 23 or September 23, 2023, to March 8, 2024. SD cards and batteries were replaced six months after deployment. For sites 5 and 6, we used sound dataset initially obtained for long-term frog monitoring on Iriomote Island (K. Fukuyama, unpublished). Recordings were scheduled with a built-in timer for 15 minutes starting at 21:00 every night in MP3 format at 128 kbps.

**Fig. 1.**
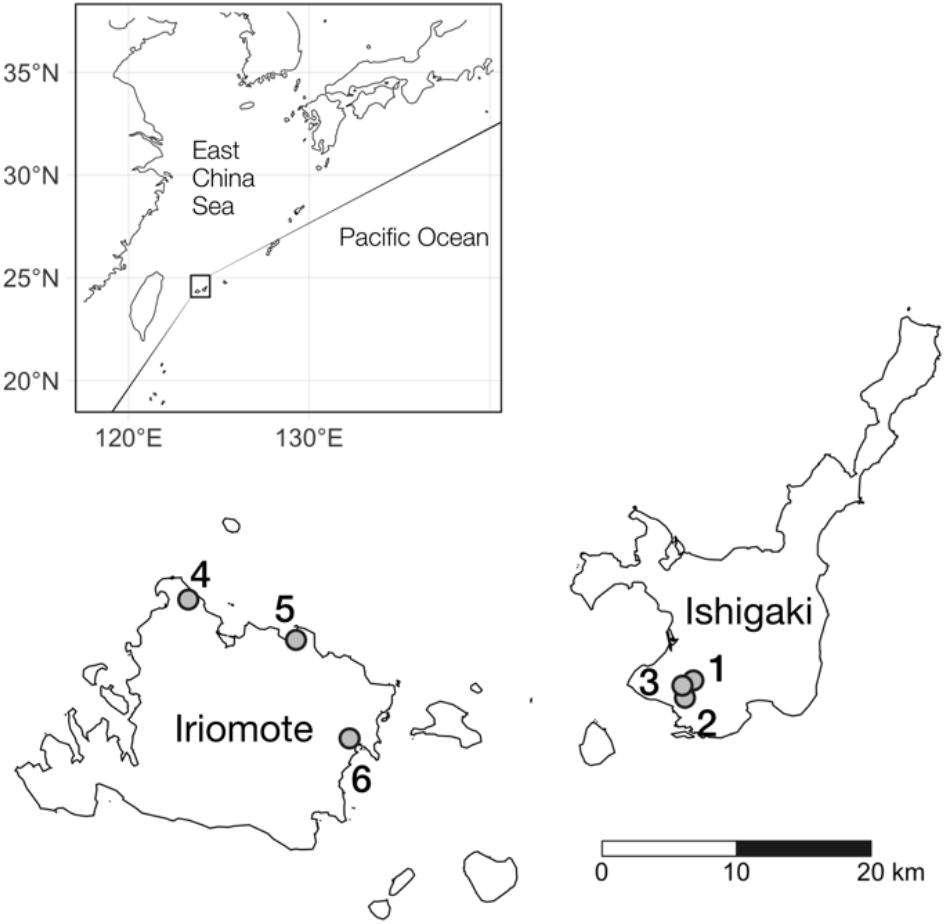
Map of study sites. Points indicate recording sites. Two alien frog species, *Polypedates leucomystax* and *Rhinella marina*, are widespread on Ishigaki Is. and have been transported to nearby islands including Iriomote Is. Map is modified from digital national land information (MLIT, Japan) (https://nlftp.mlit.go.jp)

To evaluate model performance on Iriomote Island, where the target alien frogs had no established populations, we obtained the field recordings of the alien frogs by playing back their calls. We asked a park ranger to conduct a playback survey monthly at site 4, sometime between 21:00 to 21:15. The playback sound of *P. leucomystax* was collected on Okinawajima Island and that of *R. marina* was collected on Ishigaki Island (available on the website of Ministry of the Environment, Japan: https://kyushu.env.go.jp/okinawa/wildlife/mat/m_2_2.html). The playback duration varied day by day but mostly around 60 seconds (range 30–80) for each species.

### Training and test datasets

We annotated wild and playback sounds using Deep Audio Segmenter software (Steinfath et al. 2021). For wild sounds, we selected 30 to 38 recordings per sites, totaling 190 minutes, to represent various seasons, weather conditions, and vocalizing species. These recordings were inspected through sounds and spectrograms, and we identified the beginning and end of species-specific calls following Cañas et al. (2023). Faint calls hardly visible on spectrograms were not annotated, and calls with inter-call intervals less than 0.5 seconds were combined. The annotated recordings were segmented into 3-sec audio clips to fit the model analysis unit. These audio clips were randomly split to training, validation, and test datasets in an 8:1:1 ratio. The training dataset was used to train the model, the validation dataset was for model hyperparameter tuning, and the test dataset was to evaluate model performance. The same annotation procedure was applied to the playback recordings from site 4, creating a playback sound test dataset. The playback sound dataset consists of 15.3 minutes of *P. leucomystax* calls and 12.5 minutes of *R. marina* calls. Note that no playback sounds were included in the training data.

We identified six native (*Buergeria choui, Fejervarya sakishimensis, Kurixalus eiffingeri, Microhyla kuramotoi, Nidirana okinavana, Zhangixalus owstoni*) and two alien frog species (*Polypedates leucomystax, Rhinella marina*) during the annotation procedure. For *P. leucomystax*, at least three types of calls were observed, but only the most common and conspicuous call type was annotated.

### Model training and evaluation

We used BirdNET V2.4 for custom classification model training (Kahl et al. 2021). BirdNET V2.4 is an EfficientNetB0-based classifier optimized for animal sounds (Ghani et al. 2023). While the original BirdNET is trained chiefly on bird songs, it can be applied to other taxa by fine-tuning. We used the autotune method for Bayesian hyperparameter tuning (see BirdNET documents for details: https://github.com/kahst/BirdNET-Analyzer). Number of training epochs was set to 100, and number of autotune trials was set to 200.

Model classification performance was evaluated on the wild sound test dataset and the playback sound test dataset. The model output (i.e., confidence value) more than 0.5 was considered as detection. Metrics used were precision = *TP*⁄(*TP* +*FP*) and recall = *TP*⁄(*TP* +*FN*), where TP, FP, and FN represent true positives, false positives, and false negatives, respectively.

### Visualization of model embeddings

To assess the similarity between wild and playback sounds, feature embeddings of the two datasets were visualized using Uniform Manifold Approximation and Projection (UMAP) (McInnes et al. 2018. arXiv:1802.03426). Feature embeddings are vectors obtained from an intermediate layer of a machine learning model, which can be considered as a high-level representation of acoustic signals. BirdNET V2.4 produce 1024-dimensional embeddings for a given 3-sec audio clip. These embeddings of the wild and playback datasets were plotted in 2D space with the umap package (Konopka 2023) in the R environment (R Core Team 2023). Points close in the embedding space tend to be plotted closely after applying UMAP (Coenen and Pearce 2019).

### Breeding phenology

We applied the trained model to the entire recordings collected on Ishigaki Island (sites 1–3) to infer the breeding phenology of *P. leucomystax* and *R. marina*. Reproductive activity was quantified by counting the number of detections (Kimura and Sota 2023). Give that each recording was 15 minutes in length and the BirdNET analysis unit was 3 seconds, the number of detections per recording ranged from 0 (no detection) to 300 (continuous detections throughout the recording).

## Results

### Model performance on wild sounds

Automatic hyperparameter tuning resulted in the following values; 512 hidden units, dropout rate of 0.33, batch size of 8, learning rate of 0.0001, up-sampling ratio of 1, and the use of mixup data augmentation. Model precision on the wild sound test dataset ranged from 0.850 to 1, and recall ranged from 0.794 to 1, depending on species (Table 1). For *P. leucomystax*, both precision and recall were 0.987. For *R. marina*, both precision and recall were 1.

**Table 1.** Model performance evaluated on wild sound test dataset. TP, FP, FN are the number of true positives, false positives, and false negatives, respectively. Bold font are alien species and the others are native species.

| Species | TP | FP | FN | Precision | Recall |
| --- | --- | --- | --- | --- | --- |
| ***Polypedates leucomystax*** | 77 | 1 | 1 | 0.987 | 0.987 |
| ***Rhinella marina*** | 21 | 0 | 0 | 1.00 | 1.00 |
| *Buergeria choui* | 51 | 9 | 5 | 0.850 | 0.911 |
| *Fejervarya sakishimensis* | 31 | 0 | 2 | 1.00 | 0.939 |
| *Kurixalus eiffingeri* | 27 | 3 | 7 | 0.900 | 0.794 |
| *Microhyla kuramotoi* | 95 | 7 | 8 | 0.931 | 0.922 |
| *Nidirana okinavana* | 17 | 1 | 2 | 0.944 | 0.895 |
| *Zhangixalus owstoni* | 59 | 2 | 9 | 0.967 | 0.868 |

### Model performance on playback sounds

The trained model achieved 100% precision for playback sounds of both *P. leucomystax* and *R. marina* (Table 2). Recall values were 0.629 for *P. leucomystax* and 0.906 for *R. marina*.

**Table 2.** Model performance evaluated on playback sound test dataset. TP, FP, FN are the number of true positives, false positives, and false negatives, respectively.

| Species | TP | FP | FN | Precision | Recall |
| --- | --- | --- | --- | --- | --- |
| *Polypedates leucomystax* | 149 | 0 | 88 | 1.00 | 0.629 |
| *Rhinella marina* | 192 | 0 | 20 | 1.00 | 0.906 |

When applied to the site 4 on Iriomote Island, most playback survey dates were clearly identifiable from the high number of detections (Fig. 2). For *R. marina*, few false positives or false negatives were observed. For *P. leucomystax*, the model was less accurate, with some false positives and low detection rates on May 25, July 29 and December 28. On the latter two days, the number of detections was so small that it was hard to distinguish from false positives.

**Fig. 2.**
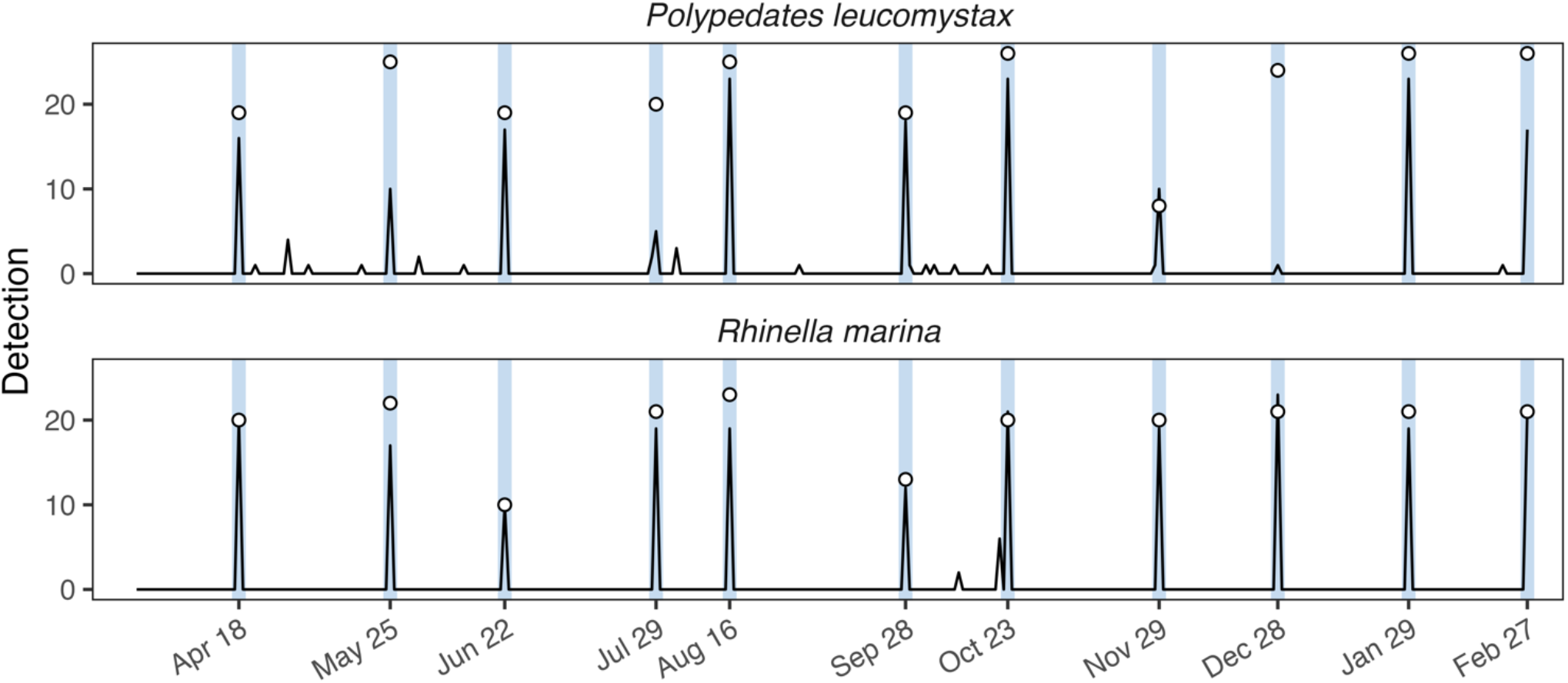
Playback call detection by the custom trained BirdNET model. Black lines are the model output, and open circles are the ground-truth values. The closer the black lines are to the white circles, the more accurate the model predictions are. Playback survey dates are highlighted in light blue.

### Visualization of model embeddings

Majority of the playback sound embeddings clustered near or overlapped with that of wild sounds (Fig. 3a). However, some audio clips containing *P. leucomystax* playback were distant from wild sound examples (Fig. 3b). On May 25 and July 29, native *Buergeria choui* called simultaneously with *P. leucomystax* (Fig. 3c). On December 28, dense choruses of *Microhyla kuramotoi* masked *P. leucomystax* playback sounds (Fig. 3d). As mentioned above, detection rates of *P. leucomystax* were low in these nights (Fig. 2). In contrast, playback sounds of *R. marina* were clearly visible on spectrograms at the same nights (Fig. 3e) and detectable by the model (Fig. 2). The results were robust to UMAP parameter settings (Supplementary Information S1).

**Fig. 3.**
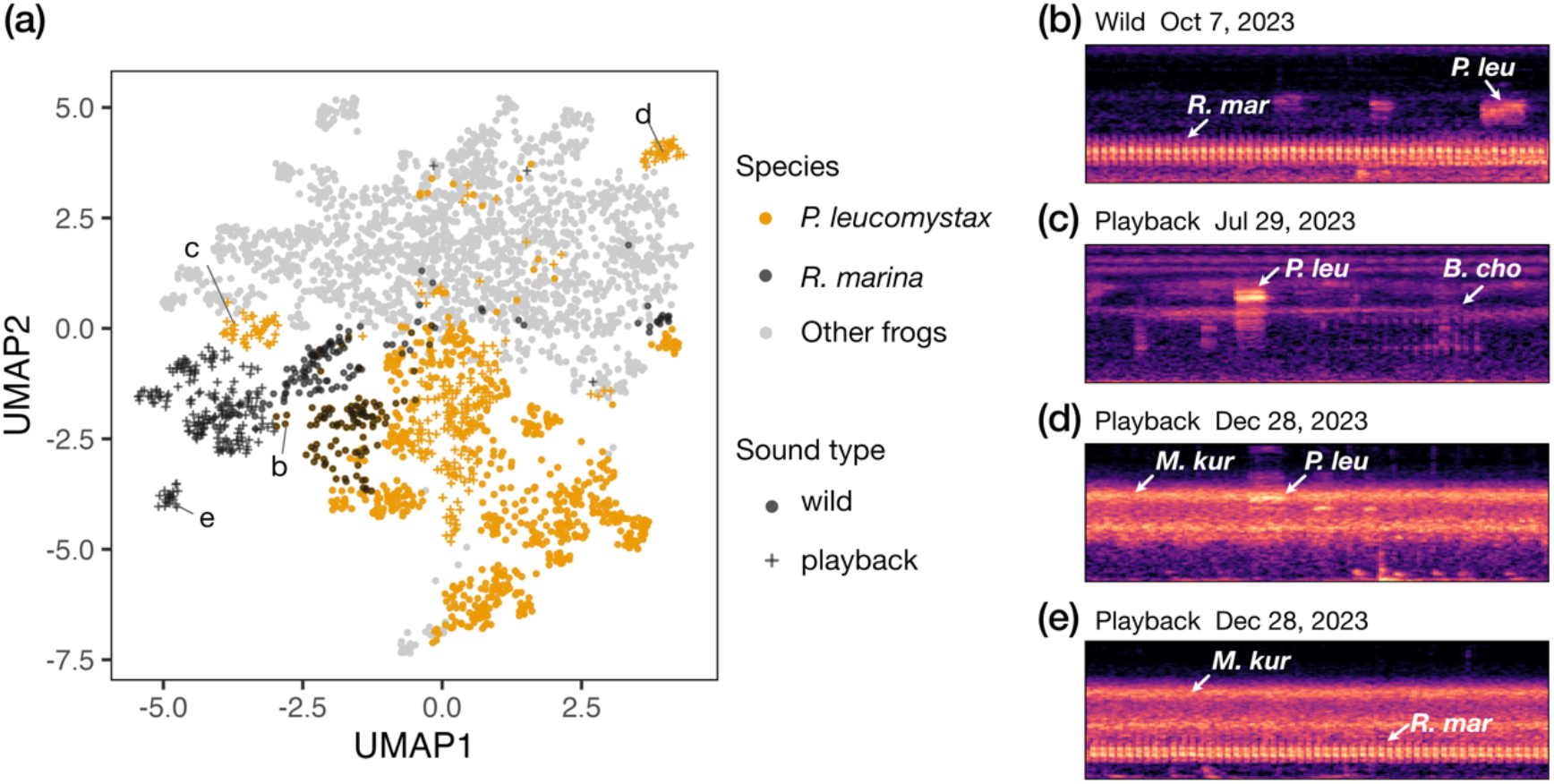
Visualization of wild and playback sound embeddings by UMAP projection (a), and examples of the corresponding mel-scaled spectrograms (b–e). Spectrograms are converted from 3 sec audio clips with the frequency range of 0 to 8000 Hz. White arrows point to some calls of *Rhinella marina* (*R. mar*), *Polypedates leucomystax* (*P. leu*), *Buergeria choui* (*B. cho*) and *Microhyla kuramotoi* (*M. kur*).

### Breeding phenology

Figure 4 shows the model’s inferences on the entire recordings on Ishigaki Island. *Polypedates leucomystax* vocalized continuously from mid-April to early November, peaking from May to September. *Rhinella marina* exhibited intermittent breeding activity without clear seasonality. Detection of *R. marina* was high in April to May 2023, September 2023, and February 2024 at site 2, and in mid-December at site 3. *Rhinella marina* was absent from site 1.

**Fig. 4.**
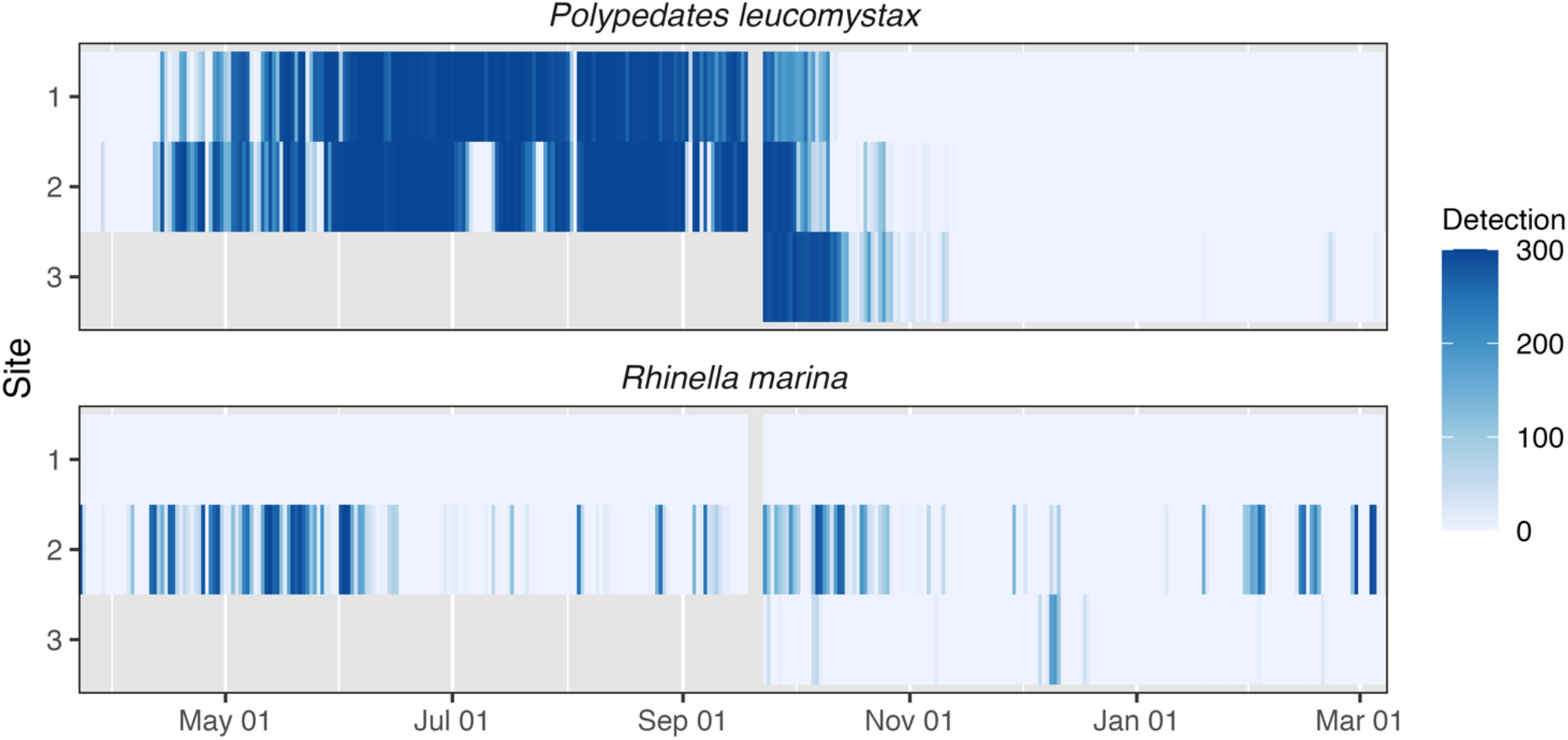
Breeding phenology of *Polypedates leucomystax* and *Rhinella marina* on Ishigaki Is. inferred from the custom trained BirdNET model. Grey color indicates missing recordings. The recording date starts from March 23, 2023, and end with March 8, 2024.

## Discussion

Using deep learning to monitor invasive alien species is increasingly popular, but its application in bioacoustics is still few (Bota et al. 2024), and the challenge of training deep learning classifiers at a pre-establishment area of invasion front has rarely been discussed. Our custom-trained BirdNET model demonstrated high accuracy not only on Ishigaki Island but also on Iriomote Island, with some reduction in recall. Despite a 9–36% decrease in recall for playback sounds compared to wild sounds (Table 2), the playback survey dates were still clearly visible in Figure 2, except on July 29 and December 28 for *P. leucomystax*. Therefore, using *ex situ* training data from Ishigaki Island did not greatly matter in our case, possibly owing to the faunal similarity between the two islands and the inclusion of training samples from both locations. Importantly, the playback survey was conducted monthly, with calls only lasting 30–80 seconds for each species in this study. But the alien frogs would vocalize more frequently in breeding seasons, and hence be easier to detect if they were introduced.

Our model has several potential applications for managing invasive species. First, it can efficiently screen passive acoustic monitoring (PAM) data for early detection of *P. leucomystax* and *R. marina*. Using PAM could reduce the cost and manpower of current manual surveillance, allowing for expansion of the survey area on Iriomote Island as well as surrounding islands that are difficult to survey frequently (e.g., Hatomajima Island; Ota et al. 2004). Second, it can inform effective surveillance schedules based on breeding phenology (Fig. 4). Third, if *P. leucomystax* and *R. marina* were introduced and established on Iriomote Island, the model could analyze PAM data to monitor its spread using site-occupancy models (Wood et al. 2020) and to quantify the impacts on native frog communities (Taylor et al. 2017).

Our workflow for training and evaluating a deep learning model is simple and can be adapted to different locations and taxa by replacing the annotated datasets. Both *P. leucomystax* and *R. marina* have been introduced to other tropical regions (Invasive Species Specialist Group ISSG 2015) and possibly continue to expand their range. Given the significant impacts of *R. marina* on local biodiversity known from studies in Australia (Shine 2010), their spread should be carefully monitored at invasion fronts. Other sound-emitting alien species such as birds, insects, and mammals can also be detected through PAM (Ribeiro et al. 2022). Deep learning-based acoustic monitoring can aid early detection of these species.

An obvious limitation of our approach is that playback calls of the target species may not perfectly mimic the soundscape at their early invasion phase. Visualization of model embeddings and the reduction of model performance suggested that the wild and playback sounds have similar features in many cases but not identical (Fig. 3, Table 2). Differences between the wild calls on Ishigaki Island and the playback sounds on Iriomote Island can be attributed to site-specific background noise in some part, but also to the playback sound property itself. Although the model effectively detected the playback sounds, similar accuracy is not guaranteed during actual applications.

Inspection of the model predictions indicated that the most common mistakes were false negatives due to faint calls. Target species can call at distance, and non-target sounds can mask them (e.g., Fig. 3d). Other studies using deep learning for PAM also listed faint calls as a common error source (Stowell et al. 2019; Bota et al. 2024). Other mistakes included misclassification of *M. kuramotoi* as *P. leucomystax*. In contrast, false positives of *R. marina* were minimal (Tables 1 and 2), likely due to the distinctively low frequency range and low variability of their calls. Adding more diverse training samples could increase model performance.

This study is the first to report the breeding phenology of *R. marina* throughout a year in Japan. Our passive acoustic survey detected their calls in all months, with peaks in spring and late summer at site 2 and in December at site 3 (Fig. 4). Chorus activity of this species is stimulated by rainfall and moderate temperature (Brodie et al. 2021). Additionally, water management of rice paddies likely influences frog reproductive activity at site 2, which was surrounded by the mixture of sugar cane fields and rice paddies. For *P. leucomystax*, the reproductive phenology on Ishigaki Island was similar to that on Okinawajima Island, another introduced population in Japan (Tanaka 2021). This species seems to be reproductively inactive during winter in Japan, unlike in Singapore (Berry 1964). These findings suggest that monitoring during spring to fall is sufficient for detecting *P. leucomystax* in this region, while year-round surveillance potentially enable early detection of *R. marina*.

## Data availability

Code and data to produce figures are available at our GitHub repository (https://github.com/kaede-kmr/Iriomote-IAS-Detect). Larger datasets for model training are deposited in Open Science Framework data repository. [to be released after the acceptance of the manuscript]

## Acknowledgements

We thank local landowners for kindly allowing us to place the acoustic recorders; Fumiko Inoue for conducting the playback survey; Chieko Matsumoto, Mitsugi Matsumoto, and the staff of Iriomote Ranger Office and Ishigaki Ranger Office for their help to conduct field surveys; Katsutoshi Watanabe for his helpful comments to our manuscript. This work was supported by ESPEC Foundation for Global Environment Research and Technology (Charitable Trust).

## Statements and Declarations

### Funding

This work was supported by ESPEC Foundation for Global Environment Research and Technology (Charitable Trust).

### Competing Interests

The authors have no relevant financial or non-financial interests to disclose.

### Author Contributions

All authors contributed to the study conception and design. Data collection was performed by Kaede Kimura and Ibuki Fukuyama. Analysis was performed by Kaede Kimura. The first draft of the manuscript was written by Kaede Kimura, and all authors commented on previous versions of the manuscript. All authors read and approved the final manuscript.

## Notes

### Competing Interest Statement

The authors have declared no competing interest.

